# When Neural Activity Fails to Reveal Causal Contributions

**DOI:** 10.1101/2023.06.06.543895

**Authors:** Kayson Fakhar, Shrey Dixit, Fatemeh Hadaeghi, Konrad P. Kording, Claus C. Hilgetag

## Abstract

Neuroscientists rely on distributed spatio-temporal patterns of neural activity to understand how neural units contribute to cognitive functions and behavior. However, the extent to which neural activity reliably indicates a unit’s causal contribution to the behavior is not well understood. To address this issue, we provide a systematic multi-site perturbation framework that captures time-varying causal contributions of elements to a collectively produced outcome. Applying our framework to intuitive toy examples and artificial neuronal networks revealed that recorded activity patterns of neural elements may not be generally informative of their causal contribution due to activity transformations within a network. Overall, our findings emphasize the limitations of inferring causal mechanisms from neural activities and offer a rigorous lesioning framework for elucidating causal neural contributions.

## Introduction

The brain is a complex network of distributed computational units whose activity profiles are assumed to reflect task-related variables, such as stimuli, or behavioral responses, such as decision variables. Decades of neuroscientific research points towards the abundance of task-relevant information in multiple spatial scales, from the coordinated activity of large-scale brain networks (*1, 2*) to the firing pattern of individual neurons (*3, 4*). For example, a line of research that started by the seminal work of Hubel and Wiesel probing into the first regions in the cat’s visual cortex (*5*) suggests that neuronal activity in the visual cortex adheres to a hierarchical organization (*6*). The firing rate of neurons in the early visual cortices was found to be higher when simple features, *e.g.,* direction and movement, were presented in stimuli, while higher-order regions are more engaged in increasingly complicated stimulus aspects such as faces and objects (*7*). Moreover, recent studies with deep artificial neural networks (ANNs) trained on images indicate a similar functional hierarchy, in which early layers encode simpler features of the input image compared to the deeper layers (*8*). These findings, supplemented by similar results from other sensory modalities such as the auditory system (*9, 10*) further provide evidence that neural networks represent features of the input in their activity profiles. An emerging line of work develops statistical and machine learning methods to reconstruct the presented input using these encoded representations (*11*).

In addition to this input-oriented view, which focuses on the relationship between the given input and the neural activity, one can study the behavior-related representations. In this case, the relationship between the activity pattern and the behavior is of interest for finding units that represent variables related to the produced behavior (*12*). To name one, the time point when a subject voluntarily moves her finger is decodable from the single cell firing pattern of neurons in the cortical supplementary motor area (*13*). Just as with the input-oriented view, emerging work develops algorithms to harness such representations, providing accurate and fast Brain Machine Interfaces (*14*). Put together, detailed analysis of neural activity patterns has proven fruitful in tackling a fundamental question of how neural networks, be they artificial or biological, encode and further use representations to produce relevant behaviors (*15*). Answering this question, however, is challenging since there is yet to be a consensus to be derived on the meaning of *representation* (*16*). More relevant to this work, it is unclear whether and to what extent recorded neural representations serve as a reliable indicator of which unit is *causally* involved in the generated behavior (*17–19*). For instance, work by Schalk and colleagues showed that the decodable information in the fusiform face area is causally irrelevant to behavior (*18*). Additionally, a recent study by Tremblay et al. found prevalent decodable task-related representations in causally irrelevant brain regions of macaque monkeys (*19*). These findings imply a dissociation between what a region represents and what it causally contributes to a given function. Said differently, it is not yet clear to what extent recorded neural representations are informative for uncovering which neural elements are causally contributing to a given function.

Other disciplines such as law (*20, 21*), economics (*22*), political, and military sciences (*23, 24*), are similarly challenged by objectively quantifying *to what extent members of a coalition are effectively contributing to a commonly produced outcome*. A game theoretical solution known as the Shapley value was proposed by Lloyd Shapley that addresses this issue (*25*). Defined simply, the Shapley value of a coalition member represents the member’s fair share of the outcome produced by the whole coalition. This value is calculated by adding the member to all possible combinations of other members and averaging the value it adds to each grouping across all these combinations (*26, 27*). To build an intuition of what the Shapley value represents, assume an orchestra in which some members contribute similarly by forming sections of similar instruments, *e.g.,* strings, while others make distinct contributions by playing solo instruments or by coordinating others, *i.e.,* the conductor (Fig 1. B). Assume that we can rearrange the orchestra by adding and removing all possible combinations of players and using an objective metric to evaluate the performed piece for every arrangement.

**Figure 1:**
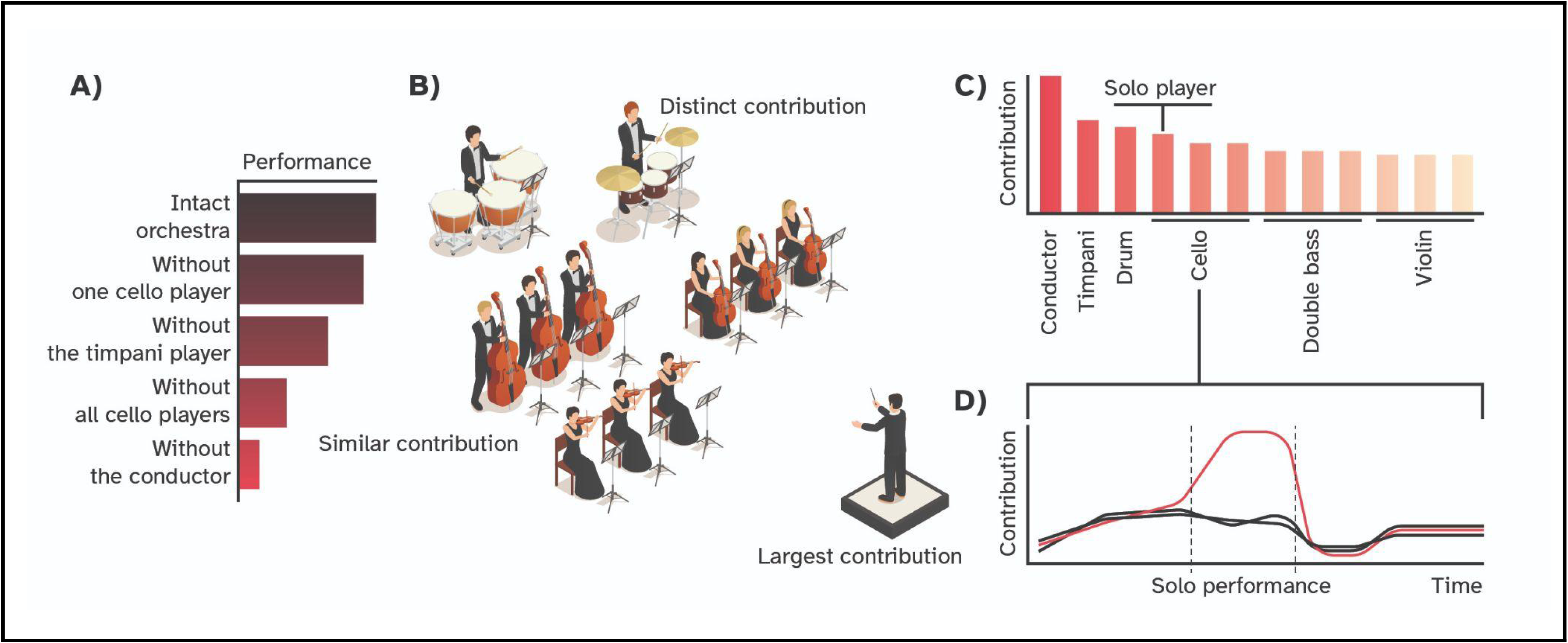
Intuition behind the notion of causal contribution. **A)** The hypothetical performance score given multiple arrangements of the orchestra. The intact network has the highest value while removing the conductor produces the largest deficit. **B)** The hypothetical orchestra in which some members have similar contributions, e.g., the string ensemble while others have distinct contributions given their role in the performed piece. **C)** Hypothetical assignment of contributions to each member given counterfactual scenarios where they were excluded from the orchestra. **D)** The hypothetical scenario where a member could have a significant contribution during a limited time period, *e.g.,* by performing a short solo piece. The isometric orchestra of the panel B is adopted from the asset designed by macrovector and is available on: freepik.com/free-vector/orchestra-isometric-composition_5967228.htm

A higher value indicates a flawless performance, while a low value indicates an unsatisfactory and erroneous performance (Fig 1. A). The Shapley value of each band member is their effective or causal contribution to this measured metric, given their contribution to every arrangement, such that the more they contribute to the performance, the larger their Shapley values would be (Fig 1. C). Intuitively, the conductor is expected to have a large Shaley value since, without him, the whole orchestra could lose its synchrony and produce many errors. Additionally, individual players in the string section are expected to have relatively smaller values, since the effect of removing one string player is usually covered by the others. However, note that the sum of their Shapley values can be large, since removing the string ensemble could also greatly impact the orchestra’s performance (Fig 1. A). Lastly, unprofessional players that introduce errors are expected to receive negative Shapley values, since removing them improves the performance. Therefore, the Shapley value represents the proportional causal contribution of each member to the produced piece.

As a substantial constraint, however, the Shapley value is not time-resolved and, as the name suggests, provides a scalar value describing each player’s contribution to the entire performance (*28, 29*). Thus, the value is insensitive to the temporal pattern in which the players contribute to the piece. For example, a cello player who is a part of the ensemble might play an outstanding short solo piece (Fig 1. D). Yet, the Shapley value assigned to her might still be mediocre overall, since other cello players in the ensemble are playing the same notes, making her role redundant for most of the performance. Moreover, the conductor’s absence could tremendously impact the orchestra at first but resolve after a while. Thus, the large initial disorganization might inflate his Shapley value. Consequently, capturing these temporal nuances provides a broader picture of effective contributions in the system, here, the orchestra. To summarize, although Shapley values represent the causal contribution of elements in a group to its outcome, they are presently static and do not provide information on the temporal profile of an element’s contribution.

In the present study, we first formalize a systematic perturbation framework to provide the objective description of which element is doing what, *i.e.,* their Shapley values, at each time point. Put differently, our method accurately captures the *time-varying causal contribution* of an element to the outcome of the coalition. Having an objective way of quantifying causal contributions of each element at each time point, we aimed to tackle the question presented above: To what extent are neural activity patterns causally informative? We first use intuitive toy examples to show how downstream nonlinear transformations applied to elements’ activity profiles can obscure their causal contributions. We then capitalize on the experimental accessibility of ANNs to perform an extensive and systematic manipulation of every combination of nodes, capturing their causal contributions (Fig. 2). Our results reveal considerable differences between nodes’ recorded activities, *i.e.,* what they represent by their activity, versus their actual causal contributions, *i.e.,* what they contribute to the produced behavior. Lastly, as with the toy examples, we show that the downstream operations applied to these neural activities could be a potential mechanism for the observed dissociation between a node’s activity pattern and its causal contribution. Altogether, this paper provides a framework for understanding when and how knowledge of neural activity may be causally irrelevant.

**Figure 2.**
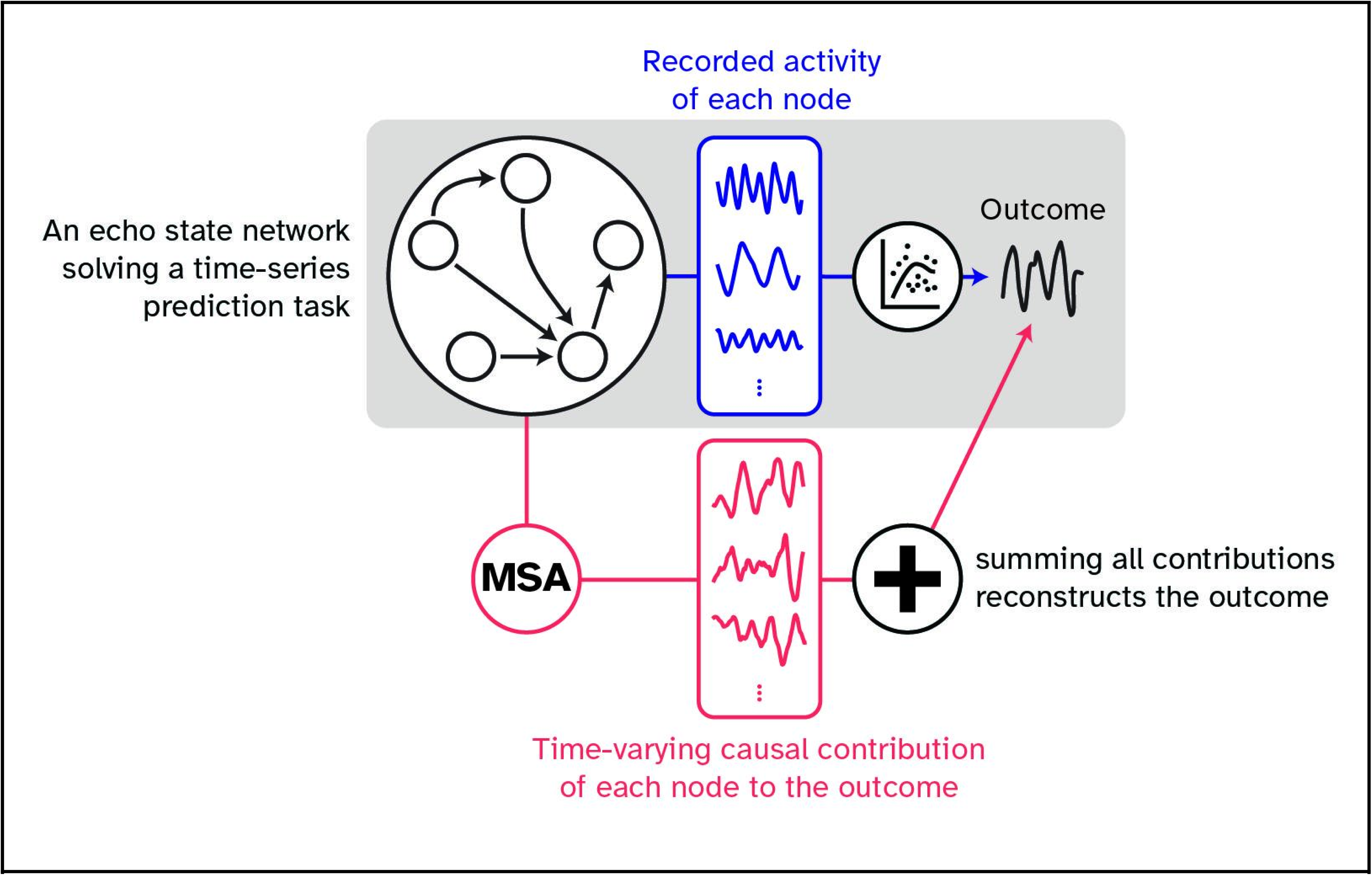
Schematic outline of the current work. In this study, we first introduced a game-theoretical framework (MSA, see Fig. 4 for more information) to precisely characterize time-varying causal contributions of elements in a given system. We then trained an ANN on a chaotic time-series prediction task and compared the nodes’ causal contributions to the network’s outcome with the nodes’ activity profiles. We found considerable differences between these two metrics, which indicates the limitations of inferring causal contributions from neural activity.

## Results

### When activity patterns are causally uninformative

In this section, we aim to provide a tangible way of differentiating what a node represents by its activity pattern and what it causally contributes to the produced outcome. To do so, it helps to first introduce an important property called ‘efficiency’ that is uniquely satisfied by Shapley value compared to other game theoretical solution concepts for allocating contributions (*25*). A solution, *i.e.,* a framework to assign causal contributions, is efficient, if the sum of contributions equals the value of the grand coalition (*30*). In other words, if the solution is efficient, then it is possible to formulate the problem starting from the outcome and consider the process as a decomposition of the outcome to its constituting building blocks. Assume three friends paid thirty dollars for a shared expense, then the efficient allocation of expenses should necessarily assign contributions that *sum up to thirty dollars*. Reformulating this example, the thirty dollars should be divided among the three friends such that no surplus or deficit is left.

To summarize the point, the causal role of a player can be defined as its *effective contribution to the outcome, such that the outcome is decomposable to these constituting contributions without any leftover surplus or deficit*. If the sum of the contributions results in a different outcome, then probably the decomposition process missed unaccounted contributions from unseen players. Moreover, note that what is meant by causal contribution depends on what is measured. For instance, we can adjust the orchestral example above by aiming to uncover *which member played what* instead of *how much they contributed to* the previously mentioned objective metric. In this case, the performed orchestral piece is decomposable to its constituting individual instruments, since it is made of nothing else but these individual instruments. Thus, it is possible to define contribution to arbitrary metrics, *e.g.,* to the ‘goodness of performance’ or the piece itself. The implications of this aspect will be further explored in the Discussion.

For the intuitive toy examples, we started by generating 30 sine-waves with various frequencies and amplitudes as independent inputs to a hypothetical neuron (Fig. 3. A). For the first example, the neuron simply summed these input activities to produce its outcome (Fig. 3. B). We then utilized multi-perturbation Shapley value analysis (MSA; Fig. 4) to derive the causal contribution of each input to the produced output signal (Fig. 3. C). Expectedly, we recovered the provided input signals simply because, in this example, the output signal was a linear summation over the inputs.

**Figure 3.**
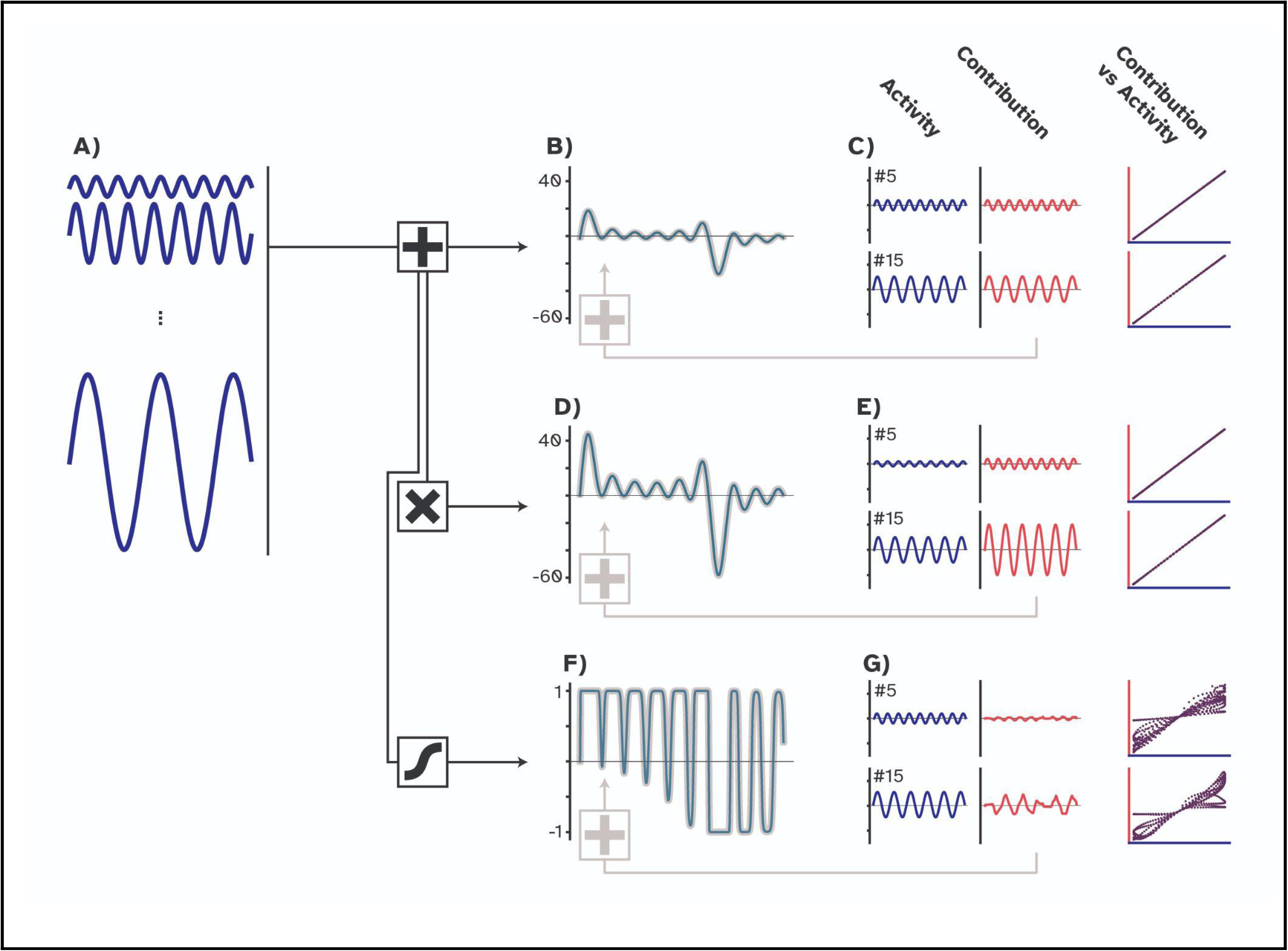
Difference between causal contributions and activity profiles in three toy examples. **A)** We first produced 30 sinusoidal waves with various frequencies and amplitudes. We called these time series building blocks. **B)** For the first example, we summed the building blocks that resulted in a new time series that we called the combined time series (teal). **C)** Employing MSA showed that the causal contributions (red) and the activity profiles (the building blocks; deep blue) are indistinguishable. Summing the contributions resulted in the combined time series (gray) since no downstream transformation was applied. **D)** For the second example, we multiplied the combined time series by a factor of two, which generated a new time series with a larger amplitude. **E)** Using MSA, we uncovered the causal contributions and found them to be twice as large as the building blocks, yet retaining a perfect linear relationship (purple), since the structure of the time series remained unchanged. Here, summing contributions resulted in the combined time series with a larger amplitude due to the downstream multiplication operation. **F)** For the last example, we passed the combined time series through a nonlinear function that resulted in a warped signal. **G)** with MSA, we found the causal contributions to be also warped. Consequently, the trajectory in the contribution-activity space is more complex. In this case, summing the contributions reconstructed the warped time series.

**Figure 4.**
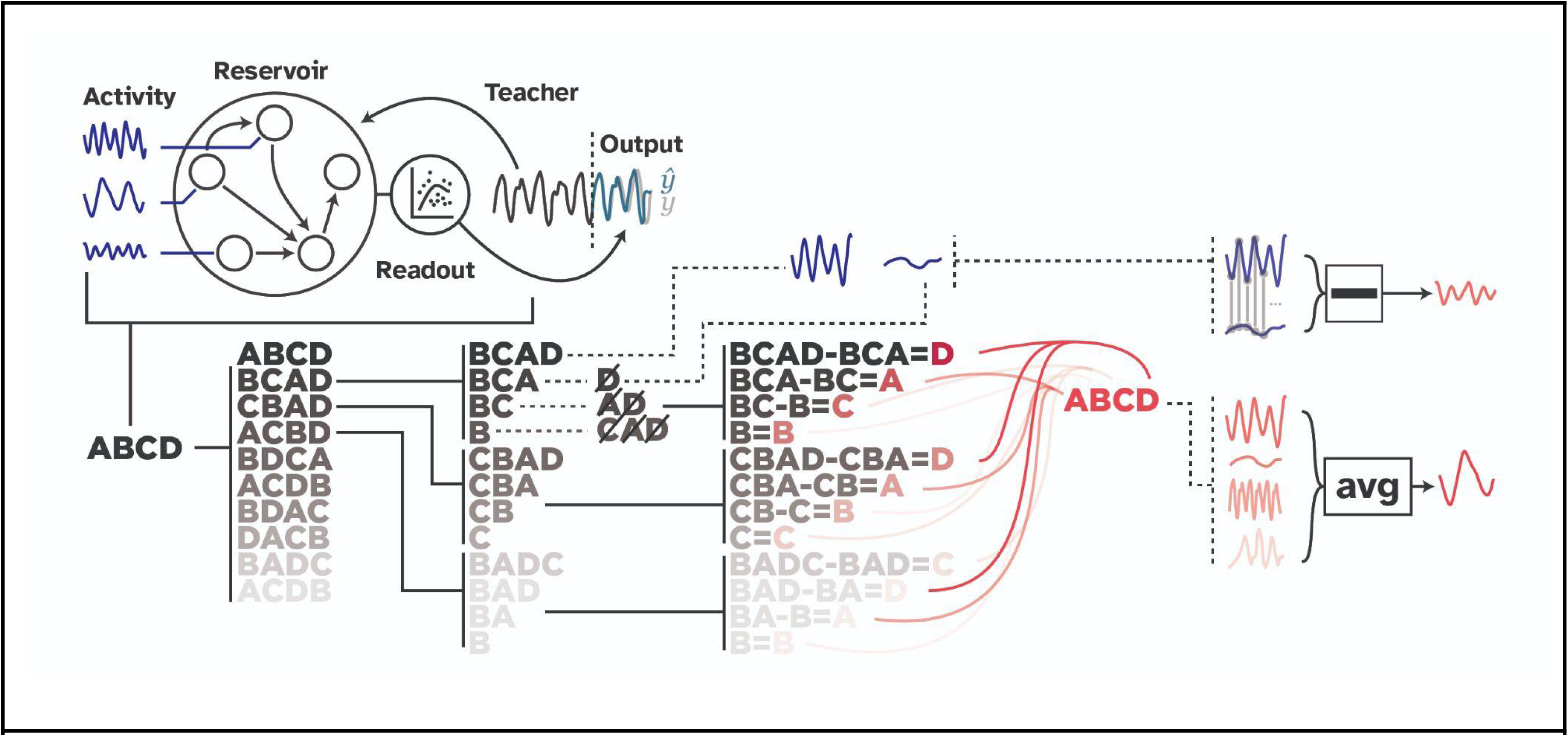
Visual abstract of the MSA algorithm and the experiment. The experimental setup is depicted on the top. The ESN was trained on a time series prediction task in which a chaotic time series was fed into the network, and a readout layer was trained on the produced representations (deep blue) of the hidden layer. After the training, the teacher sequence was disconnected, and the network was expected to generate the future time steps of the time series (teal). Note that in ESNs, the hidden layer remains unaltered during the training session. The rest of the figure depicts the MSA algorithm that was employed to compute the causal contribution of each neuron in the hidden layer. MSA samples the space of all possible combinations of node groupings to estimate nodes’ causal contributions (red). To do so, it first permutes the players and expands each permutation configuration to dictate which combinations should be perturbed. In this schematic example, to acquire the outcome produced by BCA, the example player D was removed from the game. Then to have the outcome of BC, nodes D and A were perturbed. Therefore, MSA produces a multi-site perturbation dataset that contains the outcome produced by potentially tens of thousands of unique node groupings. To then isolate the contribution of

In the second example, however, the neuron is modeled to first sum the inputs, as before, but then multiply the combined input by a factor of two (Fig. 3. D). In this case, a downstream transformation was applied to the combined inputs to produce the outcome. MSA captured the transformation, and the amplitude of the contributions was twice as large as the input signals (Fig. 3. E). Here, summing the given input resulted in the first example’s outcome, while summing the contributions resulted in the second’s (the gray thicker line in Fig. 3. B and D, respectively). The downstream multiplication applied to the input activities introduced the difference here. Note that the contributions still correlate perfectly with the input signals since each value is just doubled, and the global structure of the new outcomes remains intact. Said differently, the causal contribution of each input node to the produced outcome is not equal to the activity patterns they have produced, *i.e.,* the sine waves. The hypothetical neuron multiplied each input activity by a factor of two to produce the outcome, thus, the effective contribution to the outcome is doubled at each time point.

These two examples were intuitive, and it was easy to imagine the outcomes. To explore a non-obvious case, the third example depicts a conventional ANN neuron that combines the input and passes the result through a nonlinear hyperbolic tangent (tanh) function. In this case, the output is warped to fit within a range of -1 and 1 (Fig. 3. F). Applying MSA revealed that the contributions were also warped, and the relationship between the input and the contributions did not remain as trivial as in the previous cases. Note that the contributions were not warped as if they were separately passed through the nonlinear activation function. In other words, the order of operations matters. The transformation was applied to the *combined input* as a downstream operation, where, as observers, we have lost track of what happened to the *individual input*. MSA uncovered the impact of the downstream transformation on each input signal. Put simply, individual contributions were warped to produce the warped outcome. This conclusion is derived from the fact that the framework is efficient, and summing all contributions reconstructed the warped outcome (Fig. 3. F; gray line).

To summarize the results from this section, we first generated several sinusoidal waves as inputs to a hypothetical neuron. We showed that the only case where the input directly and fully contributed to the outcome of the hypothetical neuron was when they were simply summed to produce the outcome. We then introduced an example where a downstream transformation was applied by the neuron, here, multiplying the combined input to produce the outcome. We found that the causal contribution of each input to the output was also multiplied by a factor of two. Lastly, we introduced the common scenario in which the neuron combined the input and passed the results through a nonlinear function (tanh). We found an unintuitive impact of such downstream nonlinear transformations on the individual input signals. In the next section, we investigate how such nonlinear transformations result in warped causal contributions of neurons to the outcome of an ANN.

### Comparing representations and causal contributions in a neural network

We used a compact echo state network (ESN), that is, a class of recurrent neural networks in which the hidden layer remains unaltered during the training. We trained the network on a chaotic time series prediction task. Briefly, in this generative mode of the ESN, the network was presented with a teacher signal and had to learn a black-box model of the chaotic time-series generator. After disconnecting the teacher, the model had to produce future values of the teacher sequence (Fig. 4). Each neuron in the ESN produced representations that exhibit unique variations of the teacher signal. These representations were used by the readout mechanism to perform the task and produce the output time series (*i.e.,* the outcome of the game from a game-theoretical standpoint). Exploiting this plain model, we could investigate the impact of the transformation applied by units to other units and by the output layer to all of them. The question was posited before: *do nodes’ activity patterns readily provide causally relevant information regarding the produced output?* We hypothesized that a node’s causal contribution is a deformed version of its recorded activity, as in the nonlinear toy example. Therefore, we expected the recorded activity to be dissociated from, but still a good indicator of, causal contributions.

Employing MSA and comparing causal contributions with neural activity patterns confirmed the first part of our hypothesis, *i.e.,* the dissociation of the recorded activity from the effective causal contribution. However, as (Fig. 4. B and C) depict, the causal contribution of each node had a stark difference from its recorded activity. In other words, what a node did for the produced output signal was not trivially apparent from its activity pattern. For example, while node #32 had a faintly fluctuating activity profile, its effective contribution to the produced time series was considerably large. This means that the *almost negligible* activity of this node was highly transformed by downstream operations. Additionally, as with the toy examples, summing all causal contributions reconstructed the output signal (the thicker gray line in Fig. 5. A), indicating that MSA allocated the contributions correctly. As a sanity check, we ran the same analysis without perturbing the network and hypothesized the contributions to be zero at each time point. The rationale behind our hypothesis was the following: to compute the contributions, MSA systematically perturbs combinations of nodes and contrasts the setting where a target node was perturbed versus where it was not (Fig. 4). The difference is then assigned to be the node’s contribution to that particular group of intact nodes. If during this contrasting step, no node is removed, then the contributions of nodes are not isolated. In other words, the contrast will be between two groupings of the same configuration with no difference. The results have validated our hypothesis, and none of the nodes had any contribution to the produced output. (Supplementary Fig. 3).

**Figure 5.**
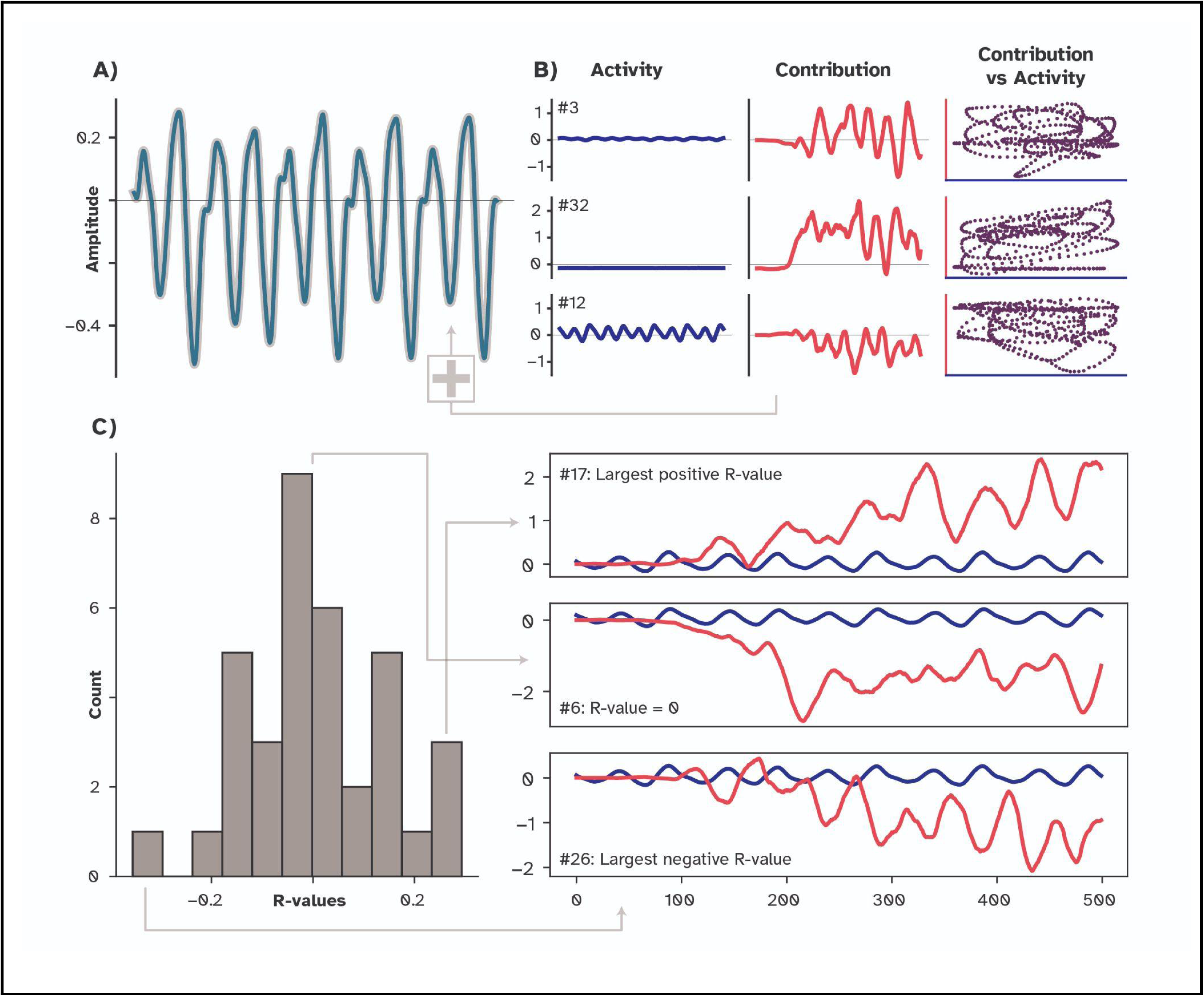
Difference between causal contributions and representations in the ESN. **A)** The line in teal shows the produced outcome (time series) by the network. Summing the uncovered causal contributions (red in panel B) perfectly reconstructs this outcome. **B)** Activity profiles (deep blue) of three exemplar nodes and their corresponding causal contributions are plotted for comparison. Note that node #32 is not completely silent but has a small range of [-0.0003, 0.007] that, relative to its causal contribution, is visually negligible. Due to the large downstream operations applied by other nodes and the readout layer, the relationship between causal contributions and neural representations (purple) is far less trivial compared to our linear toy examples (see the same plot in Fig. 4). **C)** Histogram of the Pearson’s correlation coefficient between every node’s activity profile and its causal contribution. On the right side, three examples with the largest positive, largest negative, and no correlation are plotted. Note that the activity profiles are very similar across these three nodes, while their contributions are far more diverse.

**Figure 6.**
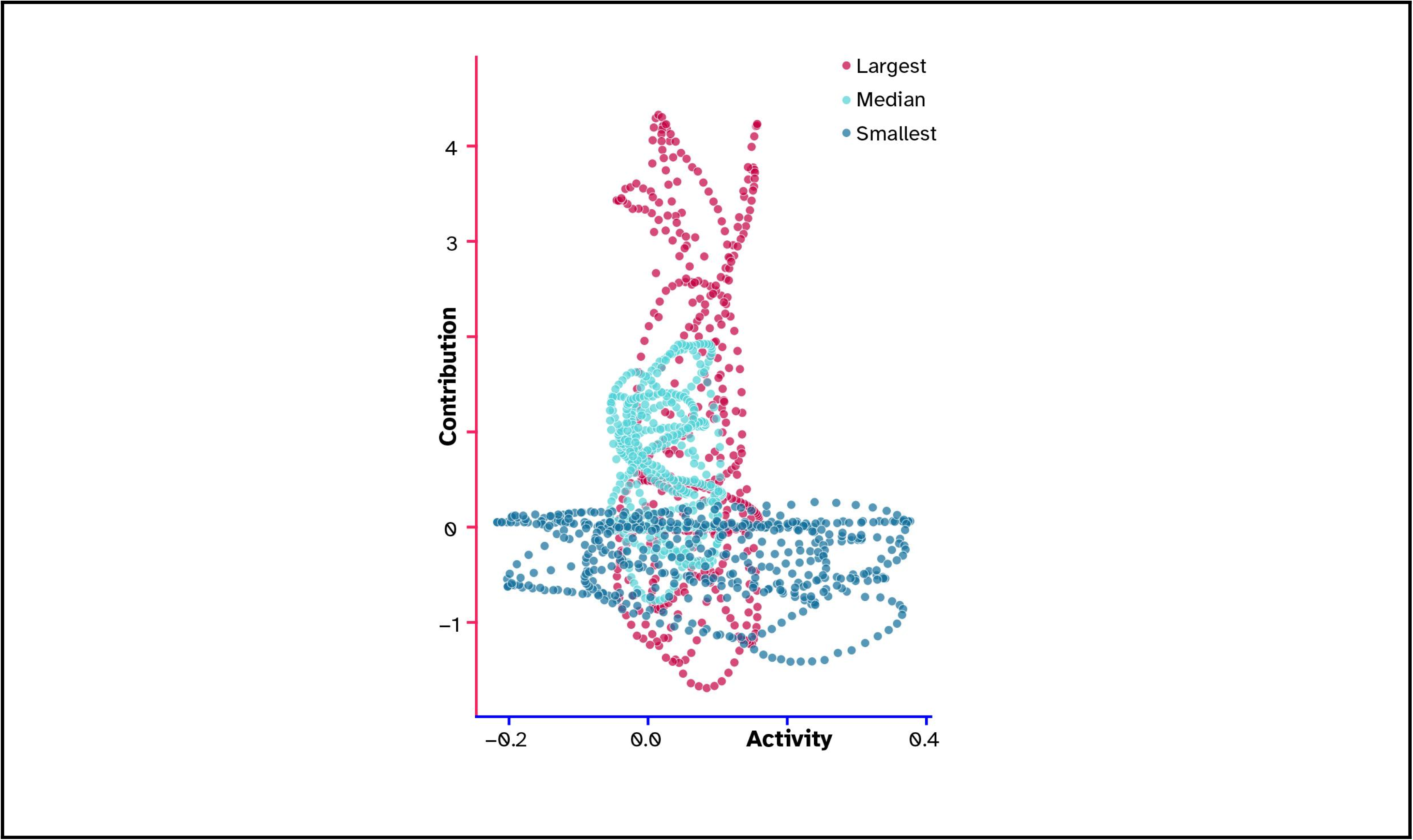
The relationship between readout weights and the transformation of representations. The trajectory of three nodes in the contribution-activity space is plotted with respect to their assigned readout weights. The node with the largest weight is plotted in red, and the one with the smallest is plotted in dark blue. The light blue depicts the same relationship for a node that was assigned the median value of the weight distribution. Note that the primary axis of variation is switched from causal contributions for the red to activity for the dark blue, while the light blue is comparatively more circular.

So far, we have shown a dissociation between the nodes’ activity profiles and their causal contributions to the produced outcome signal. With the toy examples, we showed that the downstream nonlinear operations were responsible for this observed dissociation. Our ESN example also has a downstream transformation function, *i.e.,* the readout layer. The readout layer is a regression model with a weight assigned to each neuron of the hidden layer. To see if the assigned weight can explain the observed relationship between nodes’ recorded representations and their causal contributions, we plotted this relationship for three cases: the smallest absolute readout weight, the largest absolute weight, and one in between (the median of the weight distribution). As depicted in Fig. 5, the activity pattern of the node with the largest readout weight (node #22; in red) had a relatively small range, while its causal contribution had a large variation, forming a stretched point cloud laying on the y-axis. Interestingly, this relationship had the opposite trend for the node with the smallest weight (node #12; deep blue). The point cloud was stretched on the x-axis since, relative to node #22, its contributions had a smaller variation while its activity profile had a larger variation. Lastly, the node with a weight in between also had a round and comparatively less stretched point cloud between these two extremes. We confirmed the robustness of this finding by running the analysis 50 times with different random seeds, which resulted in the same pattern (Supplementary Fig. 1). This finding suggests a relationship between the weights assigned to the readout layer (trained by fitting a linear regression model) and the casual contribution of each node to the output, such that large weights had scaled the representations more. Note that this finding can be seen as a mixture of the second and third toy examples. In the second example, the contribution was stretched due to a scaling operation (here, by the readout weights), and in the third, they were warped by the tanh operation (here, by interactions with other nodes; Fig 3. E, G).

To summarize this section, we trained a compact ESN on a time series prediction task, having a recurrent network that produced a time series while receiving no input. We then applied MSA to uncover the causal contribution of each node to the output sequence and compared them with their corresponding recorded activity. We found that the activities were heavily deformed, and what was recorded from each node did not correspond to what the node did for producing the outcome time series, *i.e.,* the behavior. Lastly, we investigated the role of the output layer and found that, as with the toy examples, the downstream operation applied to these recorded activities could be the mechanism behind the observed dissociation. Next, we discuss our findings and their implications for neuroscience.

## Discussion

In this study, we investigated a fundamental question: to what extent are recorded neural patterns causally informative of a produced behavior? To do so, we utilized a game theoretical framework, MSA, that derives the players’ fair causal contributions from their added contributions to all possible combinations of which players form coalitions. We then used three simple toy examples to build an intuition for when and how the effective causal contribution of a player may dissociate from its recorded activity. We showed that the downstream operations applied to these activity patterns are the source of the observed dissociation, with nonlinear operations having the largest impact.

Next, we applied the same pipeline to a neural network trained for solving a chaotic time series prediction task. We found that the recorded representations from neurons of the hidden layer differed considerably from their effective causal contributions to the produced output. Lastly, we were interested to see if the strengths of the readout weights in the trained network could explain the observed trajectory in the contribution-activity space. We found that the node with the largest weight also had a relatively large variation in its contribution profile, even with a faintly fluctuating activity profile. We also found the opposite trend for the node with the smallest assigned weight. It fluctuated greatly in the activity domain, but not with its causal contribution. Put together, we showed that what neurons did *for the behavior* (*i.e.,* their causal contributions) was not necessarily apparent from what they did in the network (*i.e.,* their recorded activities). To elaborate, the network’s output, which is simply obtained from multiplying the readout weights by the activity traces of the reservoir neurons, can also be reconstructed by summing up the contributions of all neurons to the behavior, here, the generated time-series. The difference between the activity and contribution of individual nodes, therefore, derives from the change in nonlinear interactions between reservoir nodes after perturbing neuronal ensembles containing a given target node.

Our work demonstrated that having access to the output (*i.e.,* the network behavior) and the recorded activities in a simple neural network without knowing the downstream operation that produces the output does not suffice to quantify the contribution of each neuron to the behavior. As argued by Barack and Krakauer “*The transformations of representations are essential for understanding cognition but are often overlooked […] To explain how behaviour is generated, […] the transformations must be identified and their neural realizations described.*” (*31*) Therefore, ignoring downstream transformations could potentially hinder one’s attempts to understand who is doing what in artificial neural networks and, to an ever greater extent, in brains. It is important to emphasize the role of transformation here. The fact that units in artificial and biological neural networks transform information is well established (*32–34*). Such systems can solve complicated tasks, probably because of their ability to harness the encoded representations to their advantage. For example, synergistic and non-redundant information is shown to be more dominant in higher-order cortical regions in the human brain, such as the prefrontal cortex (*35*), and in the deeper layers of a multi-layer neural network (*36*). In other words, higher-order regions and deeper layers of ANNs have unique and synergistic contributions to the outcome that can be due to the transformation of information along the computational chain. In ESNs, it is shown that the performance benefits from a reasonably diverse set of representations of the external stimuli (*37*). Thus, even though the prominent role of transformations in neural computations is well accepted, it is often neglected or downplayed when connecting neural activity to behavior (*38, 39*).

It is also important to emphasize the role of the quantified metric. As briefly mentioned in the Introduction, one can measure the contribution of the elements of a system to some function that quantifies an aspect of the system’s output, for instance, the performance score (*28*), or the energy of the output signal (*40*). We also noted that we could quantify the contribution of the elements to the output itself, thus bypassing the intermediate descriptive function. This is what we did in this study by computing the contribution of each node to the produced output signal itself. The difference is subtle but crucial since one is a direct deconstruction of the produced outcome while the other is a deconstruction of the description. Employing MSA on the performance score would then provide a ranking of nodes, in which the more a node contributes to the performance, the higher the value it acquires (*29*). In that case, the contributions would sum up to *the performance of the intact network,* thus deconstructing the performance score and not the outcome itself. We believe that none of these approaches is necessarily better than the other, but they are complementary, and their advantages depend on the scientific question. One could expand our work, for instance, by ranking nodes based on their contribution to the outcome and comparing the ranking with their contributions to a metric that is based on the outcome, *e.g.,* performance. Are those nodes that contribute largely to the outcome and those that contribute largely to the given metric the same? How do different metrics capture various aspects of a node’s role in the network? For example, a study could be formulated to capture the contribution of each node in an ANN to its performance, the stability of its internal dynamics, the produced outcome itself, and the global efficiency of communication in the network. Comparing nodes’ contributions to many facets of a network would then provide a far more comprehensive picture of which node is doing what and how.

ANNs aside, neuroscience relies heavily on activity patterns to infer the functional significance of some neuronal elements for the measured behavior (*41, 42*). These datasets are becoming steadily larger and more detailed, but many are yet to capture the whole brain (*43*). Our work suggests that in such cases where the downstream operations are hidden, the functional inference should be treated with caution since it is unclear if the recorded activity remains causally relevant. Even in cases where the entire brain is under scrutiny, we believe the inference could be drawn without causal claims, since a proper causal understanding of a system requires systematic and extensive multi-site manipulations of the involved units (*28, 44*). Our method, MSA, attempts to provide such a causal understanding by performing the required extensive manipulation at the expense of a relatively high computational cost. Moreover, it is unclear how causal contributions relate to the latent population manifold, uncovered by applying dimensionality reduction techniques on large-scale neural data (*14*). This relationship requires further detailed analysis, but it is a central question that needs to be addressed since it can lead to computationally cheaper estimations of causal contributions.

Here we emphasized on one network and one task for simplicity. However, we explored a separate network solving a frequency generator task where, in contrast with the main network, the exact unfolding of the target time-series is irrelevant. Our objective was to design a tunable frequency generator that utilizes the amplitude of a step function in the input to enforce the desired frequency in the output. That is, the output of the network consists of a single sinusoidal wave, with its frequency varying across different intervals. In this network too, we observed a dissociation between units’ activity profiles and their causal contributions (supplementary Fig. 4). However, to maintain a concise work and communicate a singular message, we made multiple exploratory analyses, including the differences in frequency components between the two signal sets, available in the accompanying Jupyter Notebook. Further investigation is necessary to systematically examine the relationship between the causal contributions of nodes and their activity profiles in both the frequency domain and time-frequency domain.

Additionally, in this work, both for our toy examples and the ESN experiments, we only applied one transformation to the recorded activities. It would be interesting to expand the current work by incorporating more complicated networks with deep architectures and tracking the amount of which the encoded representations transform as they propagate through a chain of nonlinear operations. Based on our findings in this paper, we hypothesize that the connection between representation and contribution becomes weaker with each layer until it reaches a point where the difference is maximal and roughly stable. It would be interesting to then investigate at which point each unit reaches its saturation point and why. Additionally, one can expand the current work by applying MSA on a network that solves multiple tasks and captures the contribution of nodes to each task. In this way, it is possible to disentangle when nodes are engaged in which tasks or if there are nodes that contribute dominantly to a specific task. Lastly, our network produced a vector (time-series) as its behavior, which resulted in a vector for the causal contribution of each node. It would be exciting to expand the contributions to higher dimensions. For example, what would be the causal contribution of a node in a generative network that produces a picture? In that case, each contribution will also be a two-dimensional image that, when summed up with other contributions, reconstructs the network’s output image. Would then nodes contribute to specific features of the image, such as the ears of the produced cat image, or would their contributions be something similar to eigenfaces, encoding a general scheme of the produced image?

To conclude, our work suggests that what neurons in an ANN represent might not necessarily and trivially map to what they causally contribute to the network’s functional output. Moreover, we provide a possible mechanism for when and how neural activity could become causally irrelevant. Lastly, we introduce a rigorous framework to capture precisely what neurons contribute to behavior, in a time-resolved manner.

## Materials and Methods

In this work, we mainly used two Python libraries. *Echoes* to implement our ESN (*45*) and *MSApy,* which is the Python implementation of the MSA framework (*46*). The code to reproduce the experiment and the supplementary analyses can be found in the GitHub repository below: https://github.com/kuffmode/contribution-modes

### Intuitive toy examples

To conduct our intuitive toy example, we initially generated a set of 30 inputs with the general form of *x*(*t*) = *Acos*(*ωt* + *ϕ*) . All signals have a similar phase, 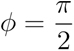. The amplitude, *A*, and frequencies, *ω* are, respectively, sampled from *A* = {0.2, 0.6, 1, 1.4, 1.8} and *W* = {1, 2.5, 4, 5.5, 7, 8.5}. Given this small dataset (see Fig. 2), we considered three linear and nonlinear output functions as 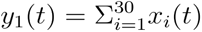, *y*_2_(*t*) = 2*y*_1_(*t*), and *y*_3_(*t*) = *tanh*(*y*_1_(*t*)).

### The echo state networks and the experimental setups

Echo state network is a multilayered artificial neural network originally proposed for learning black-box models of nonlinear systems (*37*). The core component of this model is a recurrently connected hidden layer (known as the reservoir) that receives external stimuli from an input feed-forward layer. If the connectivity profile within the reservoir is sparse, and the reservoir weight matrix fulfills certain algebraic properties in terms of singular values, internal units of this dynamical system work as the so-called “echo functions” and display unique variations of the input sequence with substantial memory contents (*47*). The echo signals are subsequently used to train a feed-forward readout layer that is the only trained part of the network. In contrast to traditional RNN training methods, where the recurrent weights in the hidden layer are gradually adapted to tune the network toward the target system, the ESN principle suggests leaving input-to-reservoir and the recurrent reservoir-to-reservoir weights unchanged after a random initialization and only optimizing the reservoir-to-output weights during training. Depending on the task, randomly generated fixed feedback connections (output-to-reservoir) may also be included in the architecture.

Given an ESN with *N_x_* leaky-integrator neurons driven by a multidimensional input signal 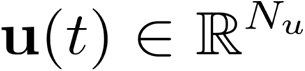, the network state update equation is described by Eq. 1,

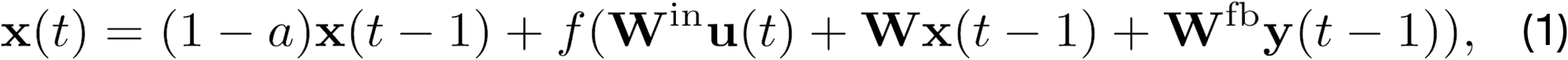

where 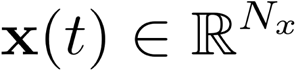 is the time-dependent *N_x_*-dimensional reservoir state and denotes the reservoir neurons’ leakage rate. The 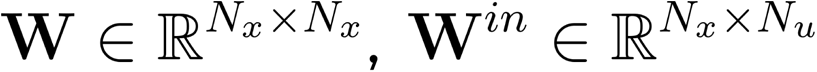, and 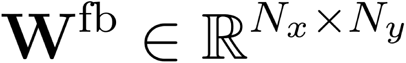 indicate internal (reservoir-to-reservoir), input-to-reservoir, and feedback (output-to-reservoir) weight matrices, respectively (*48*).

The internal states are updated via the nonlinear function, *f*, and the output 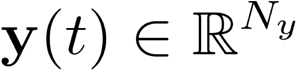 is obtained by

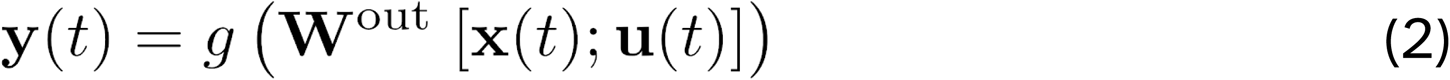

where denotes an output activation function and 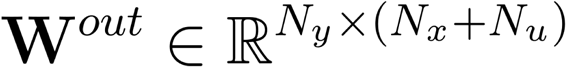 is the trainable readout weight matrix. In this equation, the extended system state, **z**(*t*) = [**x**(*t*);**u**(*t*)] is computed by the vertical vector concatenation, [.;.], of the reservoir states and external input vectors.

In this study, we conducted the ESN approach on a task of chaotic time series prediction (*37*). The objective is to learn a black-box model of the Mackey-Glass generating system that can forecast some future values of the sequence. We designed an ESN with a single neuron in input and output layers and a reservoir with a small-world topology. During the training, a fixed, low-amplitude DC signal was fed to the input as bias, and the teacher signal generated by the Mackey-Glass time-delay differential equation (Eq. 3) was presented to the output neuron.

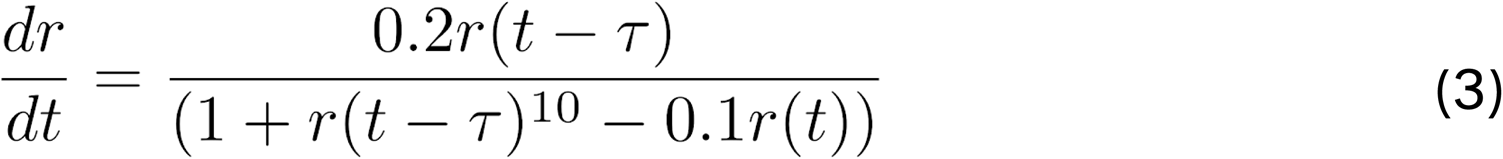

The reservoir neurons are excited by this teacher signal through the output feedback connections and, after a short transient period, start to exhibit unique nonlinear transformations of the teacher sequence. The read-out weights were computed through linear regression of the target output on these internal activation traces to minimize the least square error between the desired outputs and the network’s output signal. After training, the ESN was disconnected from the teacher and left running freely. For testing, a 500-step continuation of the original Mackey-Glass signal was computed for reference, and the mean square error between the target outputs and the network’s output was reported. Following the literature, we set the time delay *τ* = 17 and simulated Eq. 3 using the *dde2* solver for delay differential equations from the commercial toolbox MATLAB.

We used a 2500-step Mackey-Glass sequence for training and validation. The spectral radius of the internal weight matrix, *ρ*, and the neurons’ leakage rate, *α*, were optimized by performing an extensive grid search on 50 independently created 36-unit ESNs with the same small-world topology. We applied the hyperbolic tangent function, *f*(.) = *tanh*(.), for the reservoir neurons and an identity function for the readout node. Lastly, the trained network had a Mean Squared Error (MSE) of 0.0049, and the optimal spectral radius was found to be 0.66.

We conducted additional experiments on a second echo state network that was trained to perform a tunable frequency generator task. The input signal, *u*(*n*), was a slowly varying frequency setting, while the desired output, *y*(*n*), was a sine-wave of the frequency determined by the input. The input was a slow random step function, ranging from 1/25 to 1/5 Hz, and the ESN was trained on this data to produce sine-wave outputs for slow test input signals. We utilized a 2100-step input sequence for training and validation. To prevent any leakage from the previous time-point, we set the neurons’ leakage rate, a, to 1 (see Eq. 1). The hyperbolic tangent function, *f*(.) = *tanh*(.), was applied to both the reservoir neurons and the readout node. We performed an extensive grid search on 50 independently created 100-unit ESNs with random topology to optimize the spectral radius of the internal weight matrix, finding the optimal value to be 0.26. The length of the test signal was 900 steps and the trained network achieved an average MSE of 0.198 with a standard deviation of 0.036 over 500 trials. Notably, despite the differences in amplitude between the target signal and network predictions, the frequency was generated accurately at all intervals.

### Multi-perturbation Shapley value Analysis

MSA builds upon the Shapley values. The Shapley value is an axiomatic game theoretical concept that derives the fair and thus stable allocation of contributions among players of a coalition given the produced outcome by the coalition (*25*). In other words, the Shapley value of a player is its fair share of the outcome, proportional to its contribution to producing the outcome. Thus, players who contribute more will be allocated larger Shapley values, while those who do not contribute receive zero. Moreover, players who hinder the outcome receive negative values, which means that by removing them from the coalition, the outcome was increased. Theoretically, Shapley values are calculated by adding a player to all possible groupings (coalitions) of other players and tracking the outcome (*26, 29*). That is, given a coalition set,*S*, the player, *i*, and the game, *G*, with the outcome *v*, the contribution of the player to the coalition is computed as

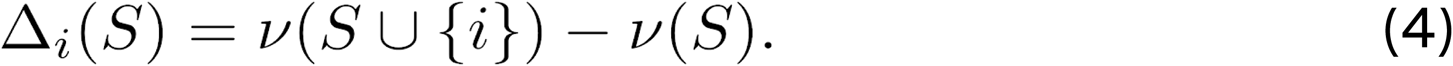

In a game with *N* players, there exists *N*! possible orderings, *R*, with length *N*. Therefore, the collective contribution of the player *i*, is defined as the average of its contribution over the set of all possible orderings, *R*, in all the coalitions to which the player is included (i.e., *S_i_*)):

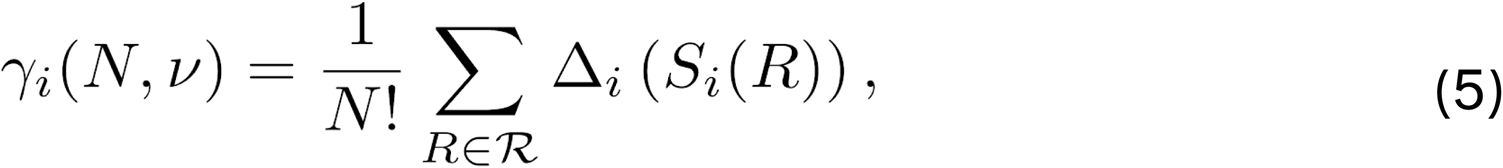

In practice, however, reaching an analytical solution to Shapley value in large sets is computationally prohibitive. Therefore, following the literature (*26, 27*), our implementation of MSA employs an unbiased estimator that randomly samples orderings from the space of possible combinations. Throughout this study, we used a sample size of *N*! that reduces the number of perturbations to *m* < *N*! unique combinations for each player and *N*^2^ × *m* in total. The MSA iterates over the produced combinations and removes the rest of the coalition to isolate the target grouping. Next, it extracts the outcome of each target grouping by playing the game. Lastly, it calculates the contribution of individual players to each of the groupings by contrasting the case where the player was in the grouping and where it was taken out alongside the rest of the removed players. After this step, the contribution of each element to each of these groupings is available. The Shapley value of each player is then its average contribution to all the sampled orderings, *R*. Algorithm 1 and Fig. 3 provide the pseudocode and a visual explanation of the MSA method.

Depending on the game, the outcome, *v*, can be either a scalar value or a vector. In this study, we simply defined the outcome as the temporal activation trace of a predefined target node in a given network, *x_i_*(*t*). To be more clear, in our toy example on *N* temporal sequences, we implemented a perturbation by simply omitting activation of the contributing input from the sum operation that produced the combined time series (see section Intuitive toy examples below). Formally, in the intact case, 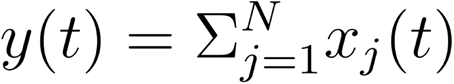, whereas for the *i*-perturbed case, 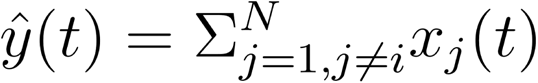. Therefore, applying Eq. 4, 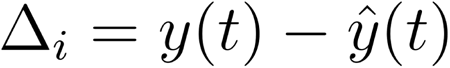.

In our ESN experiments, we trained the intact network on the reservoir neuronal activities 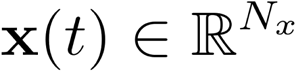 and stored the activity traces and readout weights for further analysis. We then conducted our MSA algorithm to calculate the contributions of each node to the output signal (behavior) of the intact network and to create a contribution-activity phase space as follows:

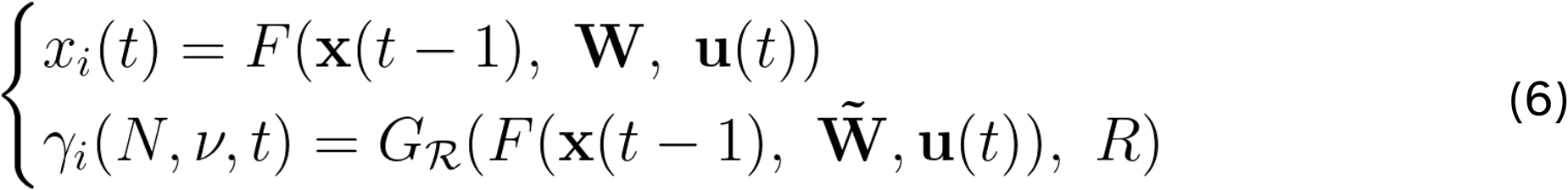

In this equation *F* represents the nonlinear operations that govern the ESN dynamics and G denotes running the ESN algorithm on the network instances with perturbed nodes. Here, W stands for reservoir weight matrix after perturbing a given node, *i*, by setting its incoming and outgoing connection weights to zero. The game, G, is played as many times as the number of all the sampled orderings, *R*. To keep the network stable after perturbations, we permitted the readout layer to be retrained after each perturbation. This allowed the network to adjust itself with the perturbed dynamics of the hidden layer as far as possible and remain functional as long as the perturbation was not severe. See the supplementary Fig. 3 for the case where this fine-tuning step was omitted. Note that the fine-tuning of the readout layer did not impact the pattern depicted in Fig. 5 since repeating the analysis with 50 different random seeds yielded the same results (Supplementary Fig. 1).

#### Algorithm 1 Multi-perturbation Shapley analysis

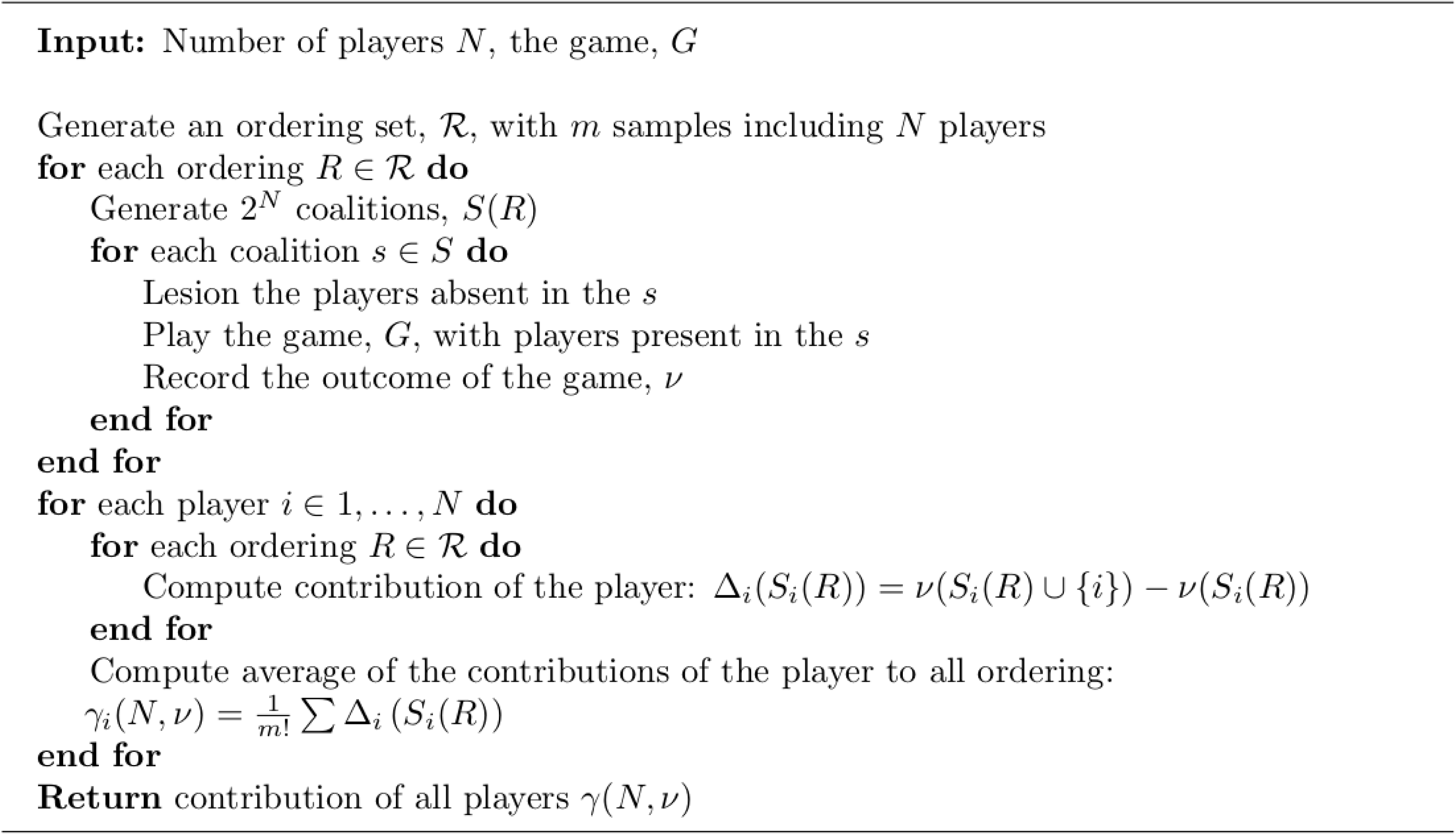

## Acknowledgments

The funding is gratefully acknowledged:

KF: German Research Foundation (DFG) - SFB 936 - 178316478-A1; TRR169-A2; SPP 2041/GO 2888/2-2.

SHD: SFB 936 - 178316478-A1. FH: DFG TRR169-A2.

KPK: NIH “Quantifying causality for neuroscience” 1R01EB028162-01.

CCH: SFB 936 - 178316478-A1; TRR169-A2; SFB 1461/A4; SPP 2041/HI 1286/7-1, the

Human Brain Project, EU (SGA2, SGA3).

The authors thank Wilhelm Braun, Fabrizio Damicelli, Mariia Popova, and Gianni Vici.

## Author contributions

Conceptualization: KF, FH, KPK, CCH. Data curation: KF, SHD.

Formal analysis: KF, FH. Funding acquisition: CCH.

Investigation: KF, SHD, FH, KPK, CCH. Methodology: KF, SHD, FH. Resources: CCH.

Supervision: FH, KPK, CCH. Validation: FH, KPK, CCH. Visualization: KF.

Writing – original draft: KF, FH.

Writing – review & editing: KF, SHD, FH, KPK, CCH.

## Data and materials availability

The code to reproduce the experiments and all relevant data is available in the GitHub repository: https://github.com/kuffmode/contribution-modes

## Supplementary Figures

**Supplementary Figure 1:**
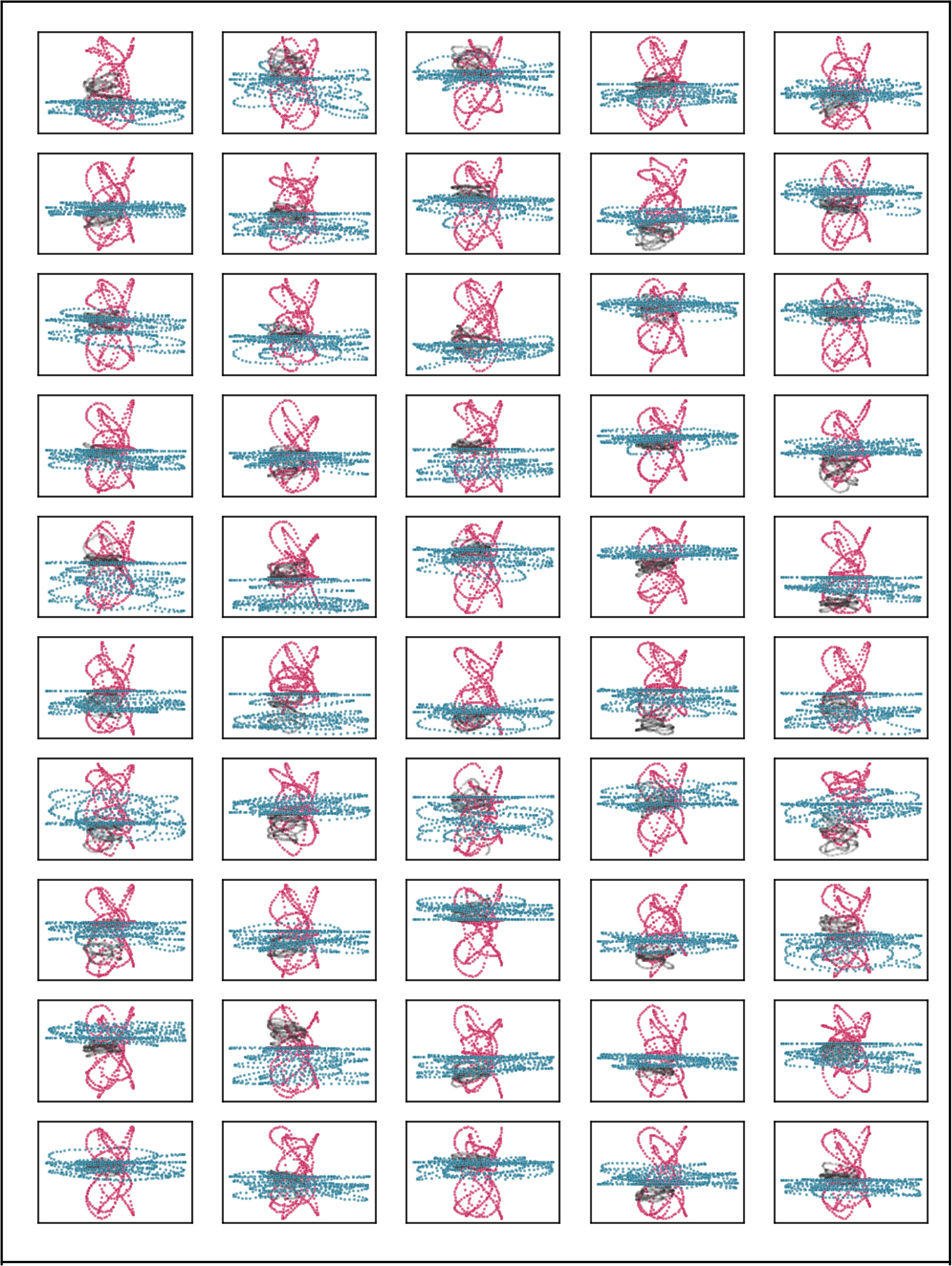
Testing with 50 different MSA random initialization. Colors and axes follow Fig. 5 with median weight plotted in black for clarity.

**Supplementary Figure 2:**
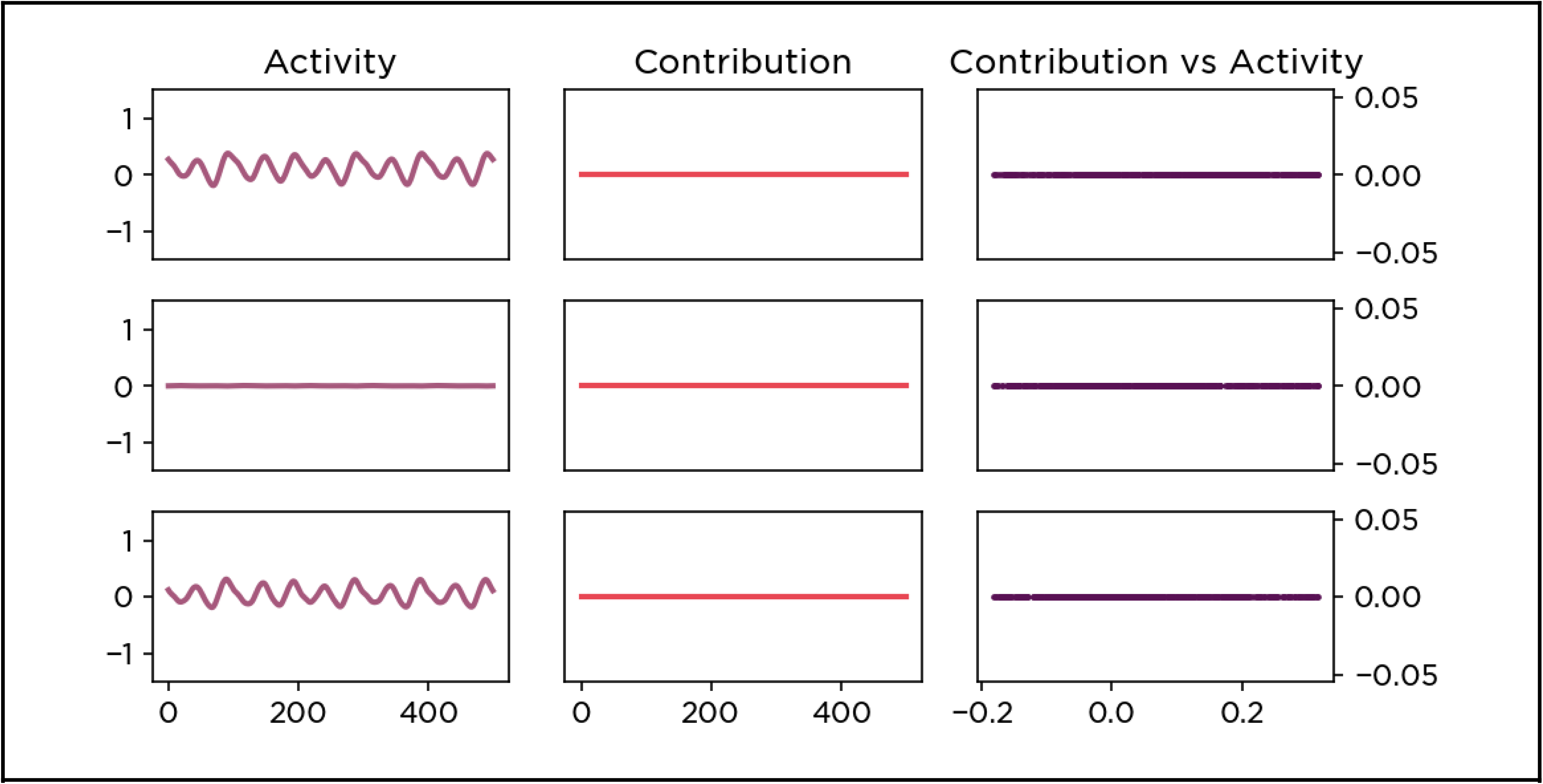
Playing games without perturbation. Activity profiles and causal contributions of the same three example nodes as in Fig.4 are plotted for comparison. Note that the contributions are all zero.

**Supplementary Figure 3:**
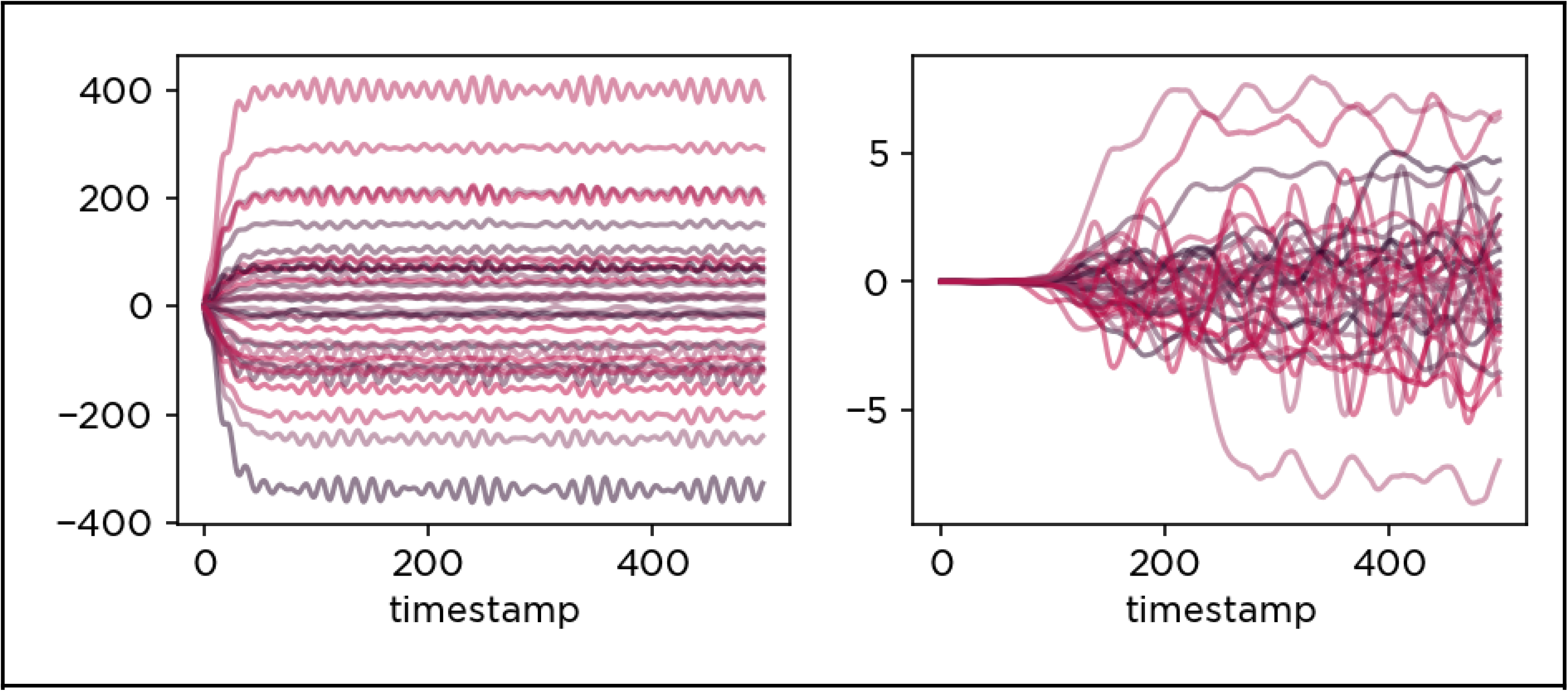
Supplementary Figure 3: Playing without retraining the readout layer. Causal contributions of all 36 nodes in the hidden layer. On the left, the fine-tuning step was omitted, and thus the network became dysfunctional given any small perturbations (note the y-axis). In right, the readout weights were allowed to adjust to the disrupted dynamics, and thus, the network could still generate a reasonable output signal.

**Supplementary Figure 4:**
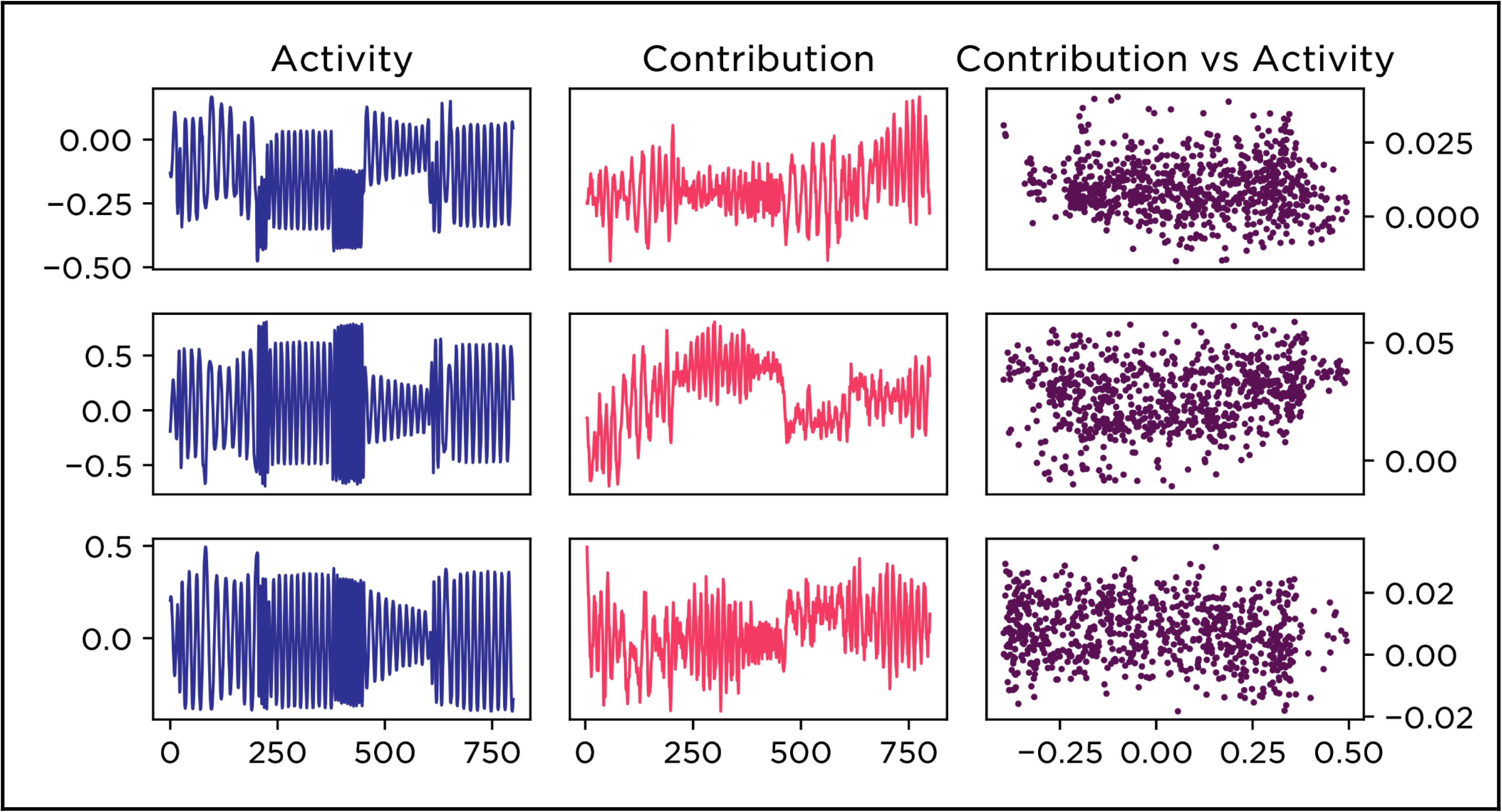
Contributions and activity profiles of a separate network solving a frequency generator task. Activity profiles and causal contributions of three example nodes in a separate network are plotted for comparison. As with the main network, the relationship between a node’s contribution and its activity profile is nontrivial.

